# Monocyte depletion attenuates the development of posttraumatic hydrocephalus and preserves white matter integrity after traumatic brain injury

**DOI:** 10.1101/388793

**Authors:** Hadijat M. Makinde, Talia B. Just, Carla M. Cuda, Nicola Bertolino, Daniele Procissi, Steven J. Schwulst

## Abstract

Monocytes are amongst the first cells recruited into the brain after traumatic brain injury (TBI). We have shown monocyte depletion 24 hours prior to TBI reduces brain edema, decreases neutrophil infiltration and improves behavioral outcomes. Additionally, both lesion and ventricle size correlate with poor neurologic outcome after TBI. Therefore, we aimed to determine the association between monocyte infiltration, lesion size, and ventricle volume. We hypothesized that monocyte depletion would attenuate lesion size, decrease ventricle enlargement, and preserve white matter in mice after TBI. C57BL/6 mice underwent pan monocyte depletion via intravenous injection of liposome-encapsulated clodronate. Control mice were injected with liposome-encapsulated PBS. TBI was induced via an open-head, controlled cortical impact. Mice were imaged using magnetic resonance imaging (MRI) at 1, 7, and 14 days post-injury to evaluate progression of lesion and to detect morphological changes associated with injury (3D T1- weighted MRI) including regional alterations in white matter patterns (multi-direction diffusion MRI). Lesion size and ventricle volume were measured using semi-automatic segmentation and active contour methods with the software program ITK-SNAP. Data was analyzed with the statistical software program PRISM. No significant effect of monocyte depletion on lesion size was detected using MRI following TBI (*p*=0.4). However, progressive ventricle enlargement following TBI was observed to be attenuated in the monocyte-depleted cohort (5.3 ± 0.9mm^3^) as compared to the sham-depleted cohort (13.2 ± 3.1mm^3^; *p*=0.02). Global white matter integrity and regional patterns were evaluated and quantified for each mouse after extracting fractional anisotropy maps from the multi-direction diffusion-MRI data using Siemens Syngo DTI analysis package. Fractional anisotropy values were preserved in the monocyte-depleted cohort (123.0 ± 4.4mm^3^) as compared to sham-depleted mice (94.9 ± 4.6mm^3^; *p*=0.025) by 14 days post-TBI. The MRI derived data suggests that monocyte depletion at the time of injury may be a novel therapeutic strategy in the treatment of TBI. Furthermore, non-invasive longitudinal imaging allows for the evaluation of both TBI progression as well as therapeutic response over the course of injury.

## Introduction

The Centers for Disease Control and Injury Prevention estimates that over 2 million people sustain a traumatic brain injury (TBI) each year in the United States, contributing to over 30% of all injury related deaths [1, 2]. In fact, TBI related healthcare expenditures near 80 billion dollars annually with an average cost of 4 million dollars per person surviving a severe TBI [3–5]. The impact of TBI is highlighted not only by its high mortality rate but also by the significant long-term complications suffered by its survivors with the progressive development of motor, cognitive, and behavioral disorders [6–10]. The immune response to TBI plays a fundamental role in the development and progression of this subsequent neurodegeneration and represents a complex interplay between peripheral immunity and the resident immune system of the injured brain [11].

Data from our laboratory, as well as others, has demonstrated that monocytes comprise a larger percentage of the early inflammatory infiltrate within the injured brain than has been formerly known [12–14]. This lead us to hypothesize that infiltrating monocytes play a more significant role in the evolution of TBI than has been previously recognized. We recently published that infiltrating monocytes contribute to the development of cerebral edema, memory loss, and locomotor dysfunction after TBI [15]. Although these infiltrating cells are short-lived within the injury milieu, we demonstrated that targeted depletion of individual circulating monocytic subsets attenuates these neurocognitive and locomotor deficits. In general, behavioral assays are effective in evaluating the effects of TBI on neurocognitive function and could be used to evaluate novel therapeutic approaches. However, behavioral data generally only allows semi-quantitative evaluation of the long-term effects of TBI progression when damage is extensive and often irreversible. Histopathological evaluation provides a direct window into the cellular and tissue alterations associated with TBI at any stage of progression. However, these types of assays require tissue harvesting and are terminal thus not allowing for the longitudinal evaluation of TBI-induced neurodegenerative processes within the same subjects. For these reasons we sought to employ a rapid and noninvasive approach to assess TBI progression and therapeutic responses following monocyte depletion over the course of TBI. Recent literature has identified ventriculomegaly, also known as posttraumatic hydrocephalus, and its associated cortical thinning as a reliable imaging marker for the progression of TBI in rodent models [16, 17]. The incidence of clinically relevant posttraumatic hydrocephalus in brain-injured human patients has been estimated be as high as 45% and the development of posttraumatic hydrocephalus is tightly correlated with clinical outcomes [18–21]. In addition, advances in neuroimaging have allowed for in-vivo assessment of traumatic disruptions between neural connections. Threshold volumetric fractional anisotropy (FA), a measure of neural connectivity, can be used to longitudinally assess the integrity of white matter pathways and disruption of functional networks after TBI [22–24]. To the best of our knowledge no study to date has applied longitudinal diffusion-weighted and 3D high resolution morphological MRI to characterize the timing and progression of TBI in an immune-depleted model of TBI. Therefore, the aim of the current study is to use a reliable and rapid MRI longitudinal assay to test whether monocyte depletion therapy can ameliorate the effects of TBI through the evaluation of 1) lesion and ventricle size and 2) white matter regional patterns associated with neuronal connectivity. We hypothesized that monocyte-depleted mice will have reduced lesion size, fewer neurodegenerative effects, and consequent preservation of normal ventricle size after TBI as compared to non-depleted controls. We also expected to observe higher values of FA across the brain in the monocyte-depleted mice as compared to non-depleted controls, demonstrating preservation of white matter and functional networks in the treated group.

## Materials and methods

### Mice

C57BL/6 male mice (n=15), 12-14 weeks old and 25g-30g, were bred and housed in a pathogen-free barrier facility within Northwestern University’s Center for Comparative Medicine. Animals were treated and cared for in accordance with the National Institutes of Health Guidelines for the Use of Laboratory Animals. The protocol was approved by the Northwestern University Institutional Animal Care and Use Committee (Protocol Number: IS00000554). All surgery was performed under ketamine and xylazine anesthesia, and post-operative analgesia was provided to minimize suffering.

### Monocyte Depletion

Mice underwent monocyte or sham depletion via intravenous injection of liposome encapsulated clodronate (Liposoma, Amsterdam) 24 hours prior to TBI as previously described [15]. Control mice were injected with liposome encapsulated PBS. Liposomes were administered according to the recommended dose of 10uL/1g of mouse. Two additional doses of liposome encapsulated clodronate, or control liposomes, were administered at 72-hour intervals post injury.

### Controlled cortical impact

Controlled cortical impact was induced as previously described by our laboratory [15]. In brief, mice were anesthetized with 50 mg/kg Ketamine (Ketaset, Fort Dodge, IA) and 2.5 mg/kg Xylazine (Anased, Shenandoah, IA) via intraperitoneal injection. A 1cm scalp incision was performed and a 5mm craniectomy was performed 2 mm left of the sagittal suture and 2 mm rostral to the coronal suture. The dura was left intact. Mice were then placed in a stereotaxic operating frame and the impactor (Leica Biosystems Inc., Buffalo Grove, IL) was maneuvered into position. A controlled cortical impact was then applied with a 3mm impacting tip at a velocity of 2.5m/s and an impacting depth of 2mm with the dwell time set at 0.1s. Immediately following injury all animals had their scalps sealed with VetBond (3M) (Santa Cruz Animal Health, Dallas, TX). All animals received post injury analgesia with Buprenorphine SR (SR Veterinary Technologies, Windsor, CO) via subcutaneous injection and were allowed to recover in separate cages over a warming pad. Mice underwent imaging at 1, 7, and 14 days post TBI and were euthanized once imaging was complete.

### MRI acquisition and image processing methods

#### MRI acquisition

Each mouse was scanned using a 7T Clinscan MRI (Bruker, Billerica, MA) using a clinical grade software platform (Syngo, Siemens) at 1, 7 and 14 days post-injury. Twenty minutes prior to scanning each mouse was injected I.P. with a dose (~0.3 mmol/Kg) of MR gadolinium based contrast agent (Prohance, Bracco) with the aim of allowing enough time for systemic delivery to the brain and leakage of agent in those regions with a breached blood brain barrier (BBB). Twenty minutes after the injection of contrast each mouse was anesthetized using isoflurane mixed with 100% O_2_ and then placed in a dedicated holder with stereotactic alignment of the head. Respiration and body temperature were monitored with the dedicated physiological monitoring system (SAI, New Jersey, USA) throughout the imaging session with anesthesia continuously delivered through a nose cone. Using a four-channel mouse brain surface coil placed on the mouse head the following sequences were used to acquire the brain images: 1) a localizer tri-axial gradient echo sequence for positioning 2) a T1-weighted gradient echo 3D sequence with TR= and TE= and isotropic spatial resolution 150 micrometers 3) a multi-direction (64 directions) and multiple values diffusion gradient (0, 200, 800) 2D EPI sequence TR= 2000msec and TE=25 msec with in plane resolution of 200 micrometers and 12 longitudinal slices covering the whole brain. Each scanning session lasted ~20-30 minutes.

#### MR Image Processing

Lesion size and ventricle volume were measured from the 3D brain images acquired at isotropic resolution of 150 micrometers using semi-automatic segmentation and active contour methods included in the freeware image processing software package ITK-SNAP http://www.itksnap.org/ [25]. The criteria for delineation of the different regions was based on the assumption that lesioned tissue (i.e. brain region affected by TBI) and ventricles exhibited a higher signal intensity than the signal from normal surrounding brain tissue (~30% higher for lesion and ~90% higher for ventricle/CSF). Two experienced preclinical imaging scientist (D.P. and N.B.) and a research scientist with experience and knowledge of brain anatomy (T.B.J.) oversaw the segmentation steps to ensure reproducibility across animals. Volume of the lesion and of the ventricular space for each mouse are expressed in mm^3^. Fractional anisotropy (FA) maps were generated using DTI processing packages contained in the image acquisition and processing platform (Syngo, Siemens) from the diffusion MRI images. In order to facilitate detection and quantification of FA differences across groups using the multi-slice data set the following approach was used. 3D rendered FA patterns were obtained using a threshold method to delineate 2D regions of higher FA in each slice using the ROI TOOL contained in the medical image processing software package JIM 7.0 (Xinapse, Essex, UK). The 3D volumetric patterns obtained with this method enabled us to quantify the global amount of white matter for each mouse by measuring the volume of brain with high FA values and to visualize in 3D abnormal connectivity patterns reflected in morphometric alterations of the rendered high FA surfaces.

### Statistical methods

The data are reported as the mean ± standard error of the mean using the statistical software program Prism (GraphPad Software Inc., La Jolla, CA). Comparison between groups for lesion size ad ventricle size was performed with two-way repeated measures ANOVA with Bonferroni’s multiple comparison posttest. Comparison between groups for fractional anisotropy were performed with a Mann-Whitney U test.

## Results

### Lesion size is unaffected by monocyte depletion after TBI

Prior studies have reported that depletion of inflammatory mediators such as neutrophils and IL-1β are able to reduce cerebral edema and prevent tissue loss after TBI [26, 27]. Our prior work has demonstrated that monocyte depletion also reduces cerebral edema and we hypothesized in the current study that monocyte depletion would reduce lesion size after TBI as well [15]. However, the prior depletion studies assessing lesion size after TBI have relied on post-mortem histopathology and gross calculation of hemispheric volume rather than specifically delineating the lesion from the ventricle enlargement that is known to occur after TBI. Furthermore, changes in lesion and ventricle size occur simultaneously making accurate characterization of the lesion more difficult. With the use of 3D, contrast-enhanced, MRI we were not able to detect significant differences in lesion size between monocyte-depleted mice (n=8) and non-depleted mice (n=7) (5 ± 1.1mm^3^ vs 7.3 ± 1.06mm^3^; *p*=0.4) (Fig 1). While this result challenges the prior literature on immune depletion and possibly suggests that monocyte depletion does not alter lesion size after TBI, it is difficult to conclude from the current data in lieu of the dynamic interplay between progressive ventricle enlargement and possible increases in lesion size, which may occur but not be detectable.

**Fig 1.**
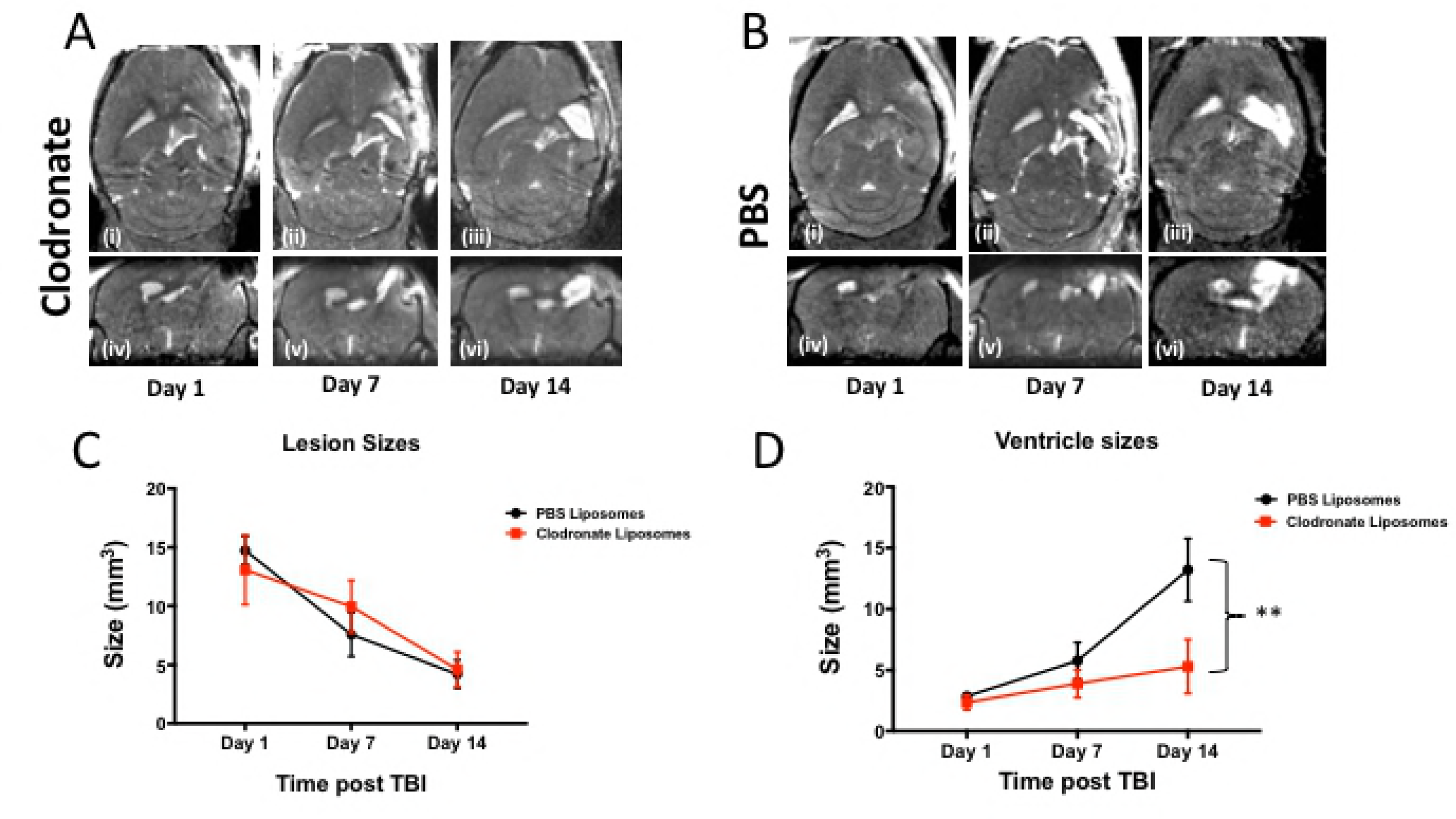
Monocyte depletion attenuates ventricle enlargement but not lesion size after traumatic brain injury. Shown are a set of representative coronal (i, ii and iii) and axial (iv, v, and vi) T1W images of a **A)** TBI/Clodronate treated mouse and a **B)** TBI/PBS treated mouse at 1-day post-TBI (i, iv), 7-days post-TBI ( ii, v) and 14-days post-TBI (iii, vi). **C)** No significant difference in lesion size was detected at any time point between monocyte-depleted and sham-depleted mice (5 ± 1.1mm^3^ vs 7.3 ± 1.06mm^3^; *p*=0.4). **D)** At 14 days post-TBI monocyte depletion markedly attenuated ventricle enlargement (5.3 ± 0.9mm^3^) as compared to sham-depleted mice(13.2 ± 3.1mm^3^; ** *p*=0.02).

## Monocyte depletion attenuates ventricle enlargement after TBI

Both lesion and ventricle size correlate with poor neurologic outcomes after TBI [28–30]. While MRI did not allow us to confirm that monocyte depletion actually reduces lesion size after TBI, we opted to determine whether monocyte depletion attenuated ventricle enlargement after TBI. While there was no difference in ventricle size immediately after injury on post-injury day 1, a non-statistically significant trend towards decreased ventricle size in monocyte-depleted mice by post-injury day 7 was measured with MRI. At 14 days post-injury monocyte depletion had markedly attenuated ventricle enlargement (5.3 ± 0.9mm^3^) as compared to sham-depleted mice (13.2 ± 3.1mm^3^; *p*=0.02) (Figs 1D and 2). These data suggest that the neurocognitive and locomotor benefits seen after monocyte depletion in our prior work may be secondary to the prevention of posttraumatic hydrocephalus after TBI [15].

**Fig 2.**
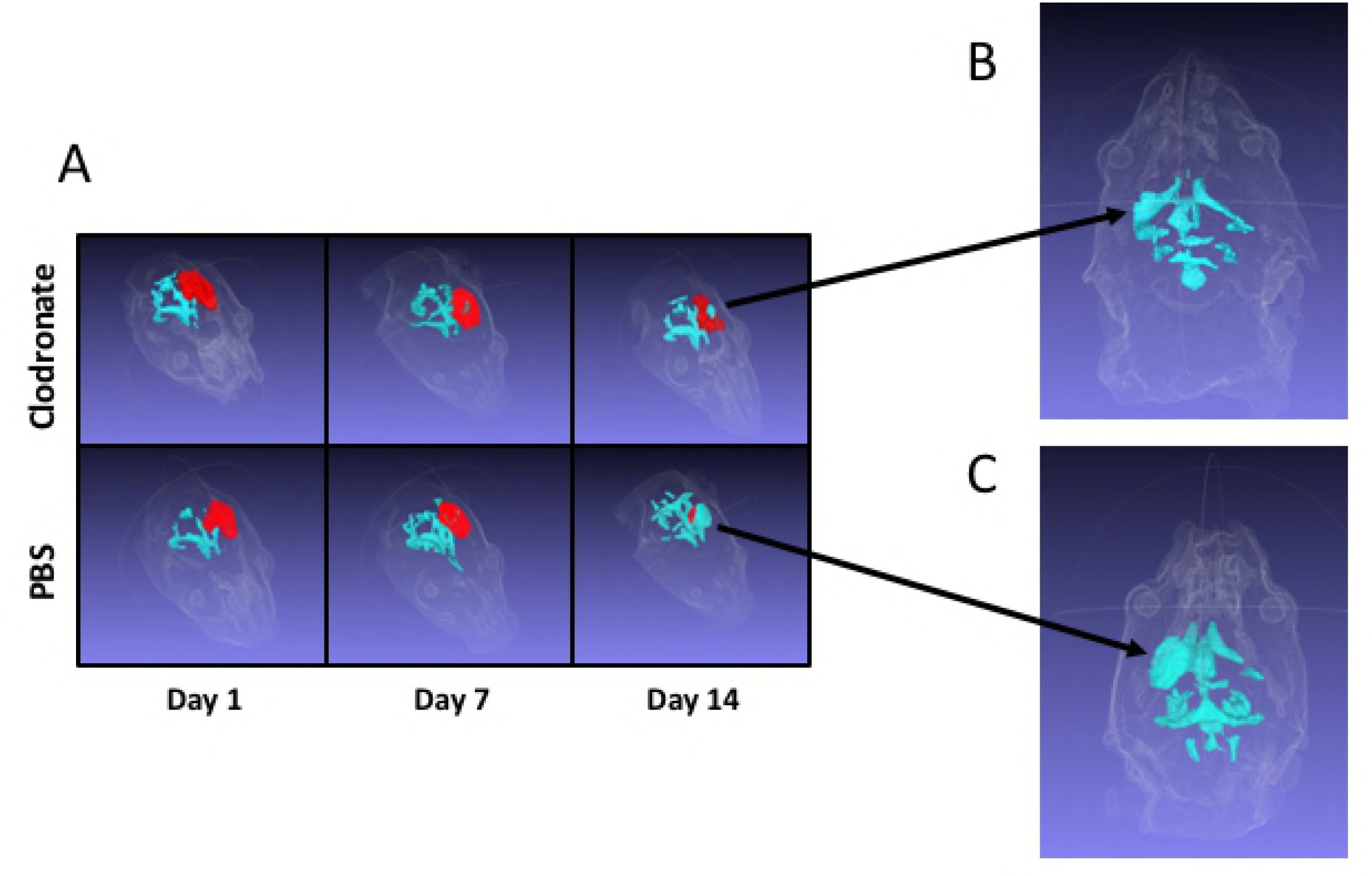
3D renderings of ventricle enlargement in monocyte-depleted vs. sham-depleted mice after traumatic brain injury. **A)** Shown in RED is the extent of traumatically injured tissue at post injury days 1, 7, and 14. Shown in light BLUE is the ventricle size. There is a progressive increase in ventricle size and decrease in injured tissue indicating loss of brain tissue and replacement with cerebrospinal fluid over the course of injury. At 14 days post-TBI the increase in ventricle size between **B)** monocyte-depleted mice (5.3 ± 0.9mm^3^) and **C)** sham-depleted mice (13.2 ± 3.1mm^3^) is markedly attenuated (*p*=0.02) indicating preservation of the injured neuronal matter.

## Monocyte depletion improves neural connectivity after TBI

Advances in neuroimaging have allowed for in-vivo assessment of traumatic disruptions between neural connections. Given that posttraumatic hydrocephalus leads to cortical thinning and loss of white matter, and that monocyte depletion prevented the development of posttraumatic hydrocephalus, we aimed to determine whether monocyte-depleted mice would show preserved white matter integrity with less disruption of functional networks after TBI. Multi-direction diffusion MRI generates images from which FA maps can be generated. The MR derived FA parameter describes restricted diffusion of water in biological tissue and in the brain has been used to effectively report on fiber density, axonal diameter, and degree of myelination in white matter [31]. Regional reduction of FA values in the brain of mice following TBI can therefore be used as a measure of white matter integrity. To better visualize global FA pattern alterations, we opted to utilize a morphometric 3D representation of FA values for each subject, generated using a threshold method as described in the methods section. This FA image analysis approach allowed us to evaluate white matter changes in the brain-injured mice from each of the two groups. As shown in Fig 3A, the volume of FA with values above a certain threshold was significantly higher in the monocyte depleted mice (123.0 ± 4.4mm^3^) at 14 days post-injury as compared to sham-depleted mice (94.9 ± 4.6mm^3^; *p*=0.025). Furthermore, both the qualitative 3D morphometric FA maps as well as the quantitative FA map exhibit extensive differences in white matter integrity and patterns between monocyte-depleted and sham-depleted mice (Figs 3B and 4). The importance of these findings lies in the ability of FA maps to predict deficits in cognitive function based the location of white matter abnormality [32].

**Fig 3.**
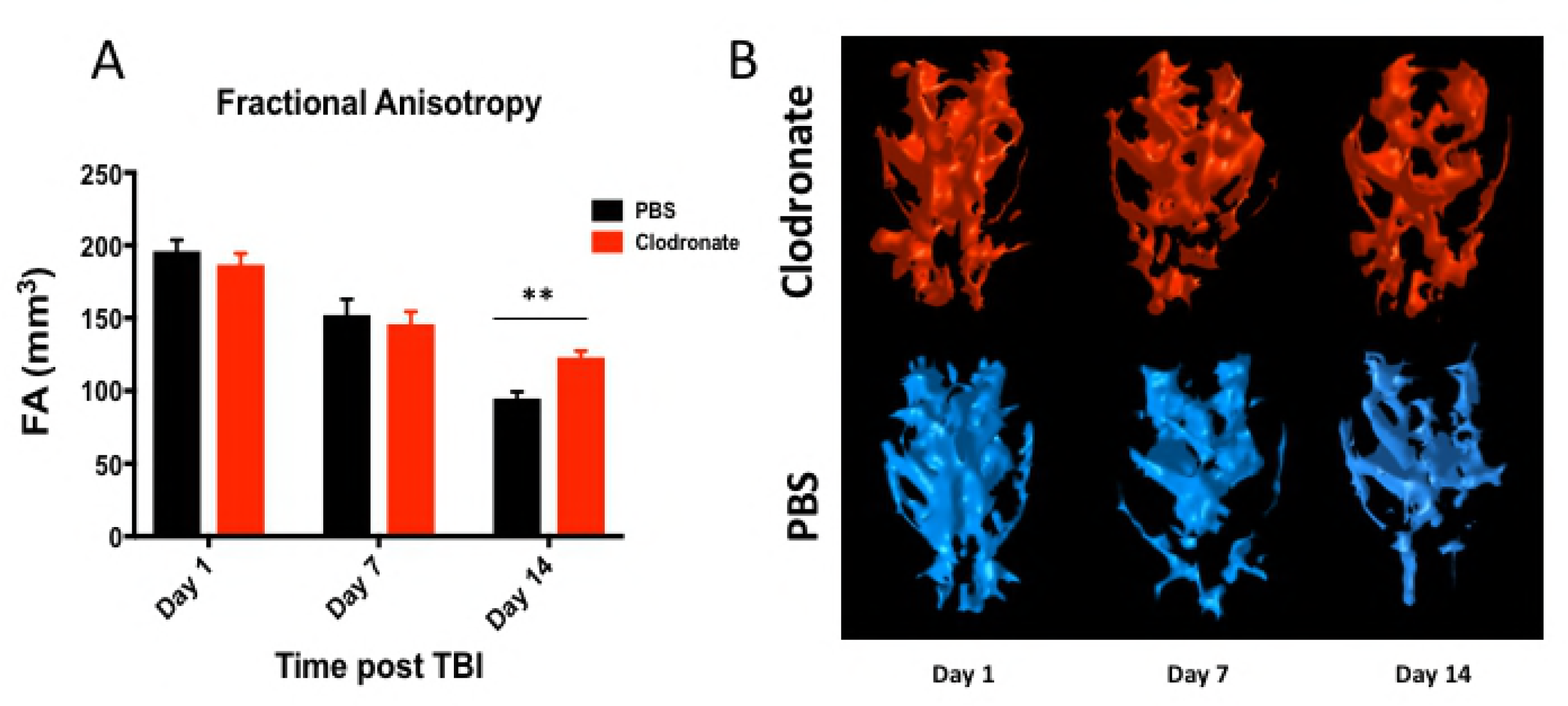
Global fractional anisotropy pattern alterations in monocyte-depleted vs. sham-depleted mice after traumatic brain injury. **A)** At 14 days post—injury the volume of fractional anisotropy above threshold was significantly higher in monocyte-depleted mice (123.0 ± 4.4mm^3^) as compared to sham depleted mice (94.9 ± 4.6mm^3^; ** *p*=0.025). **B)** Side by side 3D-rendered representative images of fractional anisotropy patterns in monocyte-depleted vs. sham-depleted mice. Noticeable is the progressive loss of pattern as the injury progresses.

## Discussion

While it is clear that the immune response to TBI plays a central role in both the resolution of the initial injury as well as in the propagation of secondary injury, the contribution of infiltrating monocytes has largely been neglected. Although, we have previously published that monocyte depletion reduces cerebral edema, improves memory, and attenuates locomotor dysfunction after TBI, to the best of our knowledge, no immune depletion study to date has deciphered in-vivo anatomic and functional outcomes during the initiation, progression, and potential resolution of TBI [15]. Magnetic resonance imaging (MRI) is a non-invasive imaging modality that can obtain structural as well as functional information in vivo. MRI utilizes low-energy radio-frequency waves in magnetic fields that allows for significantly higher resolution than ultrasound imaging or computed tomography (CT). Additionally, MRI avoids the ionizing radiation necessary for CT imaging and allows for an unlimited depth of penetration offering an advantage over the optical techniques used in histologic analysis. These advantages make MRI a valuable adjunct to behavioral and histologic assays typically used in preclinical models of disease and injury [33].

We report herein that lesion size, as assessed with contrast-enhanced 3D-MRI, was similar across both monocyte-depleted and sham-depleted mice at 1,7, and 14 days post-TBI (Fig 1A, B, and C). This indicates both a consistent and reproducible lesion across both experimental groups as well as evidence that immune modulation with monocyte depletion does not affect lesion expansion or resolution in our model of TBI. At the same time, monocyte-depleted mice demonstrated relative preservation of ventricle size at 14 days post-injury whereas sham-depleted mice developed significant posttraumatic hydrocephalus (Figs 1D and 2). This indicates that monocytes play a significant, yet poorly understood, role in the development of posttraumatic hydrocephalus. The clinical importance of this finding lies in the fact that patients who develop posttraumatic hydrocephalus demonstrate poorer recovery, require more invasive intervention, and have more complications than patients who do not develop posttraumatic hydrocephalus [18, 34, 35]. As we have shown previously, monocyte depletion abrogates neutrophil infiltration into the injured brain, improves learning, memory, motor coordination and skill acquisition [15]. We now show that monocyte depletion preserves ventricle volume after TBI thereby preventing the development of posttraumatic hydrocephalus. While the neurocognitive and locomotor benefits of monocyte depletion were detected with brain histology and neurocognitive testing, we are now able to anatomically correlate these findings in-vivo utilizing the magnetic resonance imaging techniques described in the current study.

While in-vivo MRI allows for reliable volumetric and morphometric data acquisition, it also has the ability to track anatomic and functional changes over the course of injury and disease [36–38]. Diffusion-weighted MRI uses diffusion of water within the brain to assess microstructural tissue properties. Because healthy neurons control the direction that water diffuses, diffusion imaging has the ability to map the white matter of the brain. Differential diffusion within white matter can be used to detect demyelination and axonal degeneration in the form of increased water diffusion through injured myelin structures [39]. This is quantified through fractional anisotropy (FA), and lower FA associates with white matter injury. These FA measurements allow for quantification of different properties of tissue structure and function which can be applied across experimental groups over time [40]. In the current study, our data demonstrates that monocyte-depleted mice have significantly higher FA as compared to sham-depleted mice after TBI (Fig 3A). This indicates that monocyte depletion not only attenuates the development of posttraumatic hydrocephalus, but also conserves functional white matter microstructure as shown by the preserved quantitative FA volumetrics as well as the qualitative FA maps generated after TBI (Fig 3B). These findings correlate well with our prior work demonstrating preserved working memory, better motor coordination, and improved skill acquisition in monocyte-depleted mice [15].

White matter tracts are bundles of axons connecting functionally specific, yet anatomically separated, regions of the brain. When a TBI is sustained, these axons are subjected to shear, compression, and torsion forces. Recent advances in diffusion MRI techniques, such as diffusion tensor imaging (DTI) and high angular resolution diffusion imaging (HARDI), are now allowing for damage to specific white matter bundles to be identified and correlated with neurocognitive functions that are known to associate with those bundles [41]. While the tresholded FA analysis used in the current study is similar to DTI-MRI analysis approaches that can provide fiber tractography visualization, our method has the advantage of a higher sensitivity with respect to DTI techniques. Nonetheless, efforts are underway to compare the two approaches with the aim of exploring specific connectivity patterns using fiber tractography (33). All of these techniques offer an abundance of opportunity for the detection of injuries that might otherwise go unnoticed with more conventional imaging. This has the potential to offer more individualized diagnoses that can lead to more individualized treatment in the future. However, despite the excitement surrounding these new techniques there remains several methodologic obstacles to overcome before cutting edge neuroimaging will be incorporated into patient care [42]. Perhaps the biggest challenge will be for clinicians to incorporate the data generated from these studies into clinical decision making.

**Fig 4.**
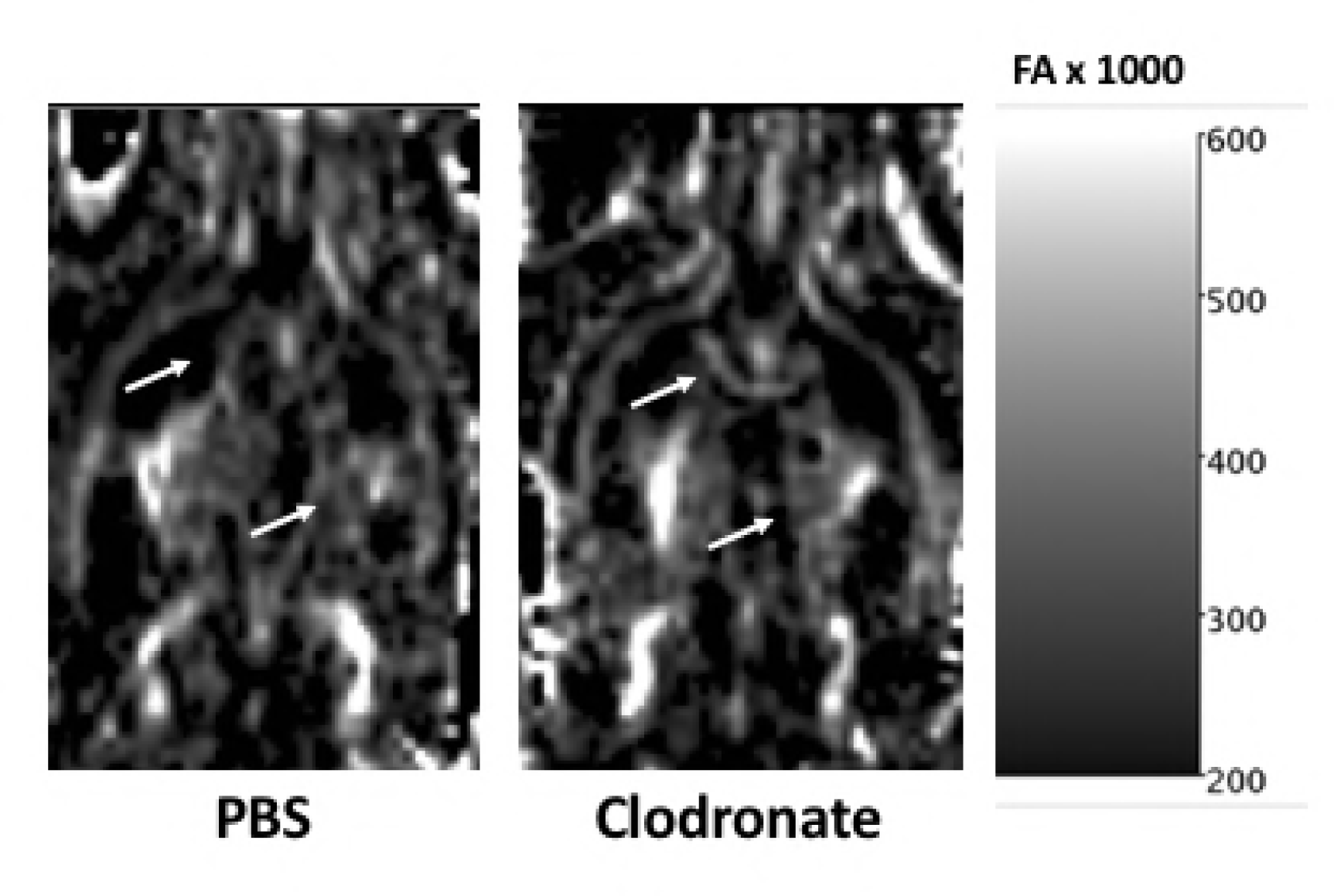
Representative quantitative fractional anisotropy (FA) map 14-days post injury. White arrows showing regions of reduced/altered FA patterns in a sham-depleted mouse as compared to a monocyte-depleted mouse.

## Conclusions

In conclusion, we report that while monocyte depletion does not reduce lesion size in mice after TBI, it does attenuate the development of posttraumatic hydrocephalus as assessed by 3D, contrast-enhanced, MRI. Furthermore, diffusion MRI and the resultant quantitative FA volumetrics and qualitative FA mapping show preservation of functional white matter tracts in monocyte-depleted mice as compared to sham-depleted mice after TBI. These data suggest that peri-injury depletion of monocytes may represent a novel therapeutic strategy for the treatment of TBI with the potential to be translated to human patients.

